# A novel crustavirus associated with tail fan necrosis in New Zealand red rock lobsters, *Jasus edwardsii*

**DOI:** 10.1101/2025.03.10.642514

**Authors:** Rebecca M. Grimwood, Leonardo N. Zamora, Jessica A Darnley, Lizenn Delisle, Kate S. Hutson, Julie Hills, Jemma L. Geoghegan

## Abstract

Tail fan necrosis (TFN) is a shell disease affecting spiny lobsters’ outer integument, with significant implications for the health and commercial viability of red rock lobsters (*Jasus edwardsii*) in New Zealand. Despite its impact, the potential role of a microbial agent in TFN remains poorly understood. Here, we conducted metatranscriptomic analyses on matching uropod and haemolymph samples from 15 red rock lobsters exhibiting TFN symptoms to characterise the associated microbial communities and search for putative candidates for further investigation. We report the discovery of a novel crustavirus (family: *Nyamiviridae*), named Red rock lobster crustavirus (RRLCV), consistently present in the uropod tissues of all sampled lobsters with TFN. RRLCV’s presence and high abundance in uropod tissues suggest a potential association with TFN, although we found no direct relationship between viral abundance and TFN severity metrics, such as tissue loss. We also identified 30 bacterial genera across uropod samples, including previously associated groups, however, these were detected inconsistently. Our findings raise new questions about the tissue tropism, transmission, and potential pathogenicity of RRLCV in red rock lobsters. While TFN appears to be a multifactorial condition, RRLCV represents a promising candidate for further research into its role in the development of this disease.

## Introduction

Crustaceans are an incredibly diverse set of arthropods and play significant roles in aquatic ecosystems (Rowley, 2022). Among these, decapod crustaceans, which include shrimp, crabs, and lobsters, are revered in global markets due to their high commercial value (Dhar et al., 2022; Francis, 2022). Consequently, the growth of decapod trade has driven increasing interest in the diseases affecting these species, particularly lobsters coming from commercial fisheries or aquaculture production (Behringer et al., 2012; Longshaw, 2011; Shields, 2011; Sindermann, 1989). While certain pathogens are well characterised (Dhar et al., 2022; de Souza Valente & Wan, 2021), our understanding of the full spectrum of their infectious diseases remains incomplete. Shell diseases that affect lobsters’ outer integument (carapace) pose significant challenges to their trade due to the impact on their marketability and consumer acceptability. Importantly, shell diseases often lack known aetiologies (Rowley & Coates, 2023).

Tail fan necrosis (TFN) is a type of classical shell disease affecting several species of spiny lobster, characterised by progressive deterioration and melanisation of the uropod and telson (Shields, 2011). A case definition was recently proposed by Jones et al. (2024) that considers population source, clinical signs and histopathological assessment. Symptoms include blistering, blackening and erosion of tail fan tissue leading to visible lesions and, in severe cases, significant tissue loss (Geddes et al., 2005; Jones et al., 2024). Shell diseases affecting crustaceans have a wide global distribution, however, syndromes involving initial blistering of the tail fan have only been observed in Australia and New Zealand (Diggles et al., 2022; Jones et al., 2024). This condition not only affects the physical appearance and commercial value of lobsters but also impacts their swimming efficiency and overall health (Musgrove et al., 2005; Rowley & Coates, 2023).

Red rock lobsters, *Jasus edwardsii*, (also known as crayfish or kõura) (Hutton, 1875) are a species of spiny lobster found in the coastal waters of New Zealand and Australia (Kensler, 1967). They are important both commercially and culturally in New Zealand (Booth, 2006; Chandrapayan et al., 2009) and are significantly impacted by TFN (Musgrove et al., 2005). The history and prevalence of TFN in New Zealand rock lobsters are not well documented, but previous reports indicate incidences of up to 17% in male lobsters in the North Island (Freeman & MacDiarmid, 2009). Furthermore, TFN has been recognised as an increasing issue within the fishing industry (Pande et al., 2021; Zha et al. 2017).

Multiple aetiologies for TFN have been suggested, though the exact cause remains under investigation. Factors such as bacterial infections, environmental stressors like overcrowding, water quality, pollution, and handling practices during capture and processing have been considered as potential contributors (Freeman & MacDiarmid, 2009; Musgrove et al., 2005, Zha et al. 2018a, 2019). Currently, no single microbial agent has been consistently associated with the syndrome, but repeated handling and damage to the tail fan followed by subsequent secondary bacterial infection has been a leading hypothesis (Geddes et al., 2004; Jones et al., 2024).

Viruses are abundant in aquatic environments (Bergh et al., 1989), but they have rarely been implicated in TFN (Pande et al., 2021). Comprehensive studies on marine invertebrate viromes are also limited, possibly resulting in undetected cases. Viral pathogens are found in other crustacean species (Bateman, 2021; Bateman & Stentiford, 2017) but are not well classified. The lack of crustacean cell lines has also made it difficult to study and characterise viruses *in vitro* (Bateman & Stentiford, 2017). Metagenomic methods, however, permit the identification and study of microbial organisms independently of prior knowledge and more traditional methods (Li et al., 2015). Still, despite invertebrates constituting the majority of the world’s animal species, gaps and biases in our knowledge surrounding their host-virus interactions (Alfonso et al., 2024) warrant more thorough investigations, especially in the face of emerging aquatic diseases with unknown causes.

Understanding and mitigating TFN is key to maintaining the health and commercial viability of New Zealand’s rock lobster populations. This study aimed to investigate the eukaryotic, prokaryotic, and viral microbial communities associated with matching uropod and haemolymph samples. A total of 15 red rock lobsters that were either suspected of exhibiting TFN when captured or had developed TFN following four weeks in captivity were subject to a metatranscriptomic investigation to identify putative candidates with a possible role in the progression of this condition.

## Methods

### Animal ethics, transport, and husbandry

All procedures performed involving live lobsters were carried out in accordance with New Zealand government regulations and associated ethical standards as administered by the Nelson Marlborough Institute of Technology Limited Animal Ethics Committee approval (Ref. AEC2023-CAW-06).

Fifteen live lobsters were sourced from licenced commercial fishers. Lobsters were captured off the northeastern region of the South Island, New Zealand (Figure 1a). After landing, the lobsters were transported to Cawthron Institute’s aquatic physical biocontainment (PC2) facility, Te Wero Aro-anamata, Nelson, New Zealand. A rapid visual health assessment to identify any obvious external damage including TFN-like symptoms was performed before the lobsters were allocated to three independent recirculating seawater aquaculture systems that included a 1,000 L circular tank (i.e. n= 5 lobsters per tank). Each tank was fitted with particle filtration, inline ultraviolet (UV) disinfection, foam fractionator and biofiltration. Natural seawater was sourced from the Cawthron Aquaculture Park and was UV-treated and filtered to 5 μm before use. The lobsters were held for up to four weeks prior to sampling, during which water quality was monitored via a supervisory real-time data acquisition system. Lobsters were fed once daily to satiation using 4.5 mm extruded pellets (Biomar Quinnat Plus IQ).

**Figure 1.**
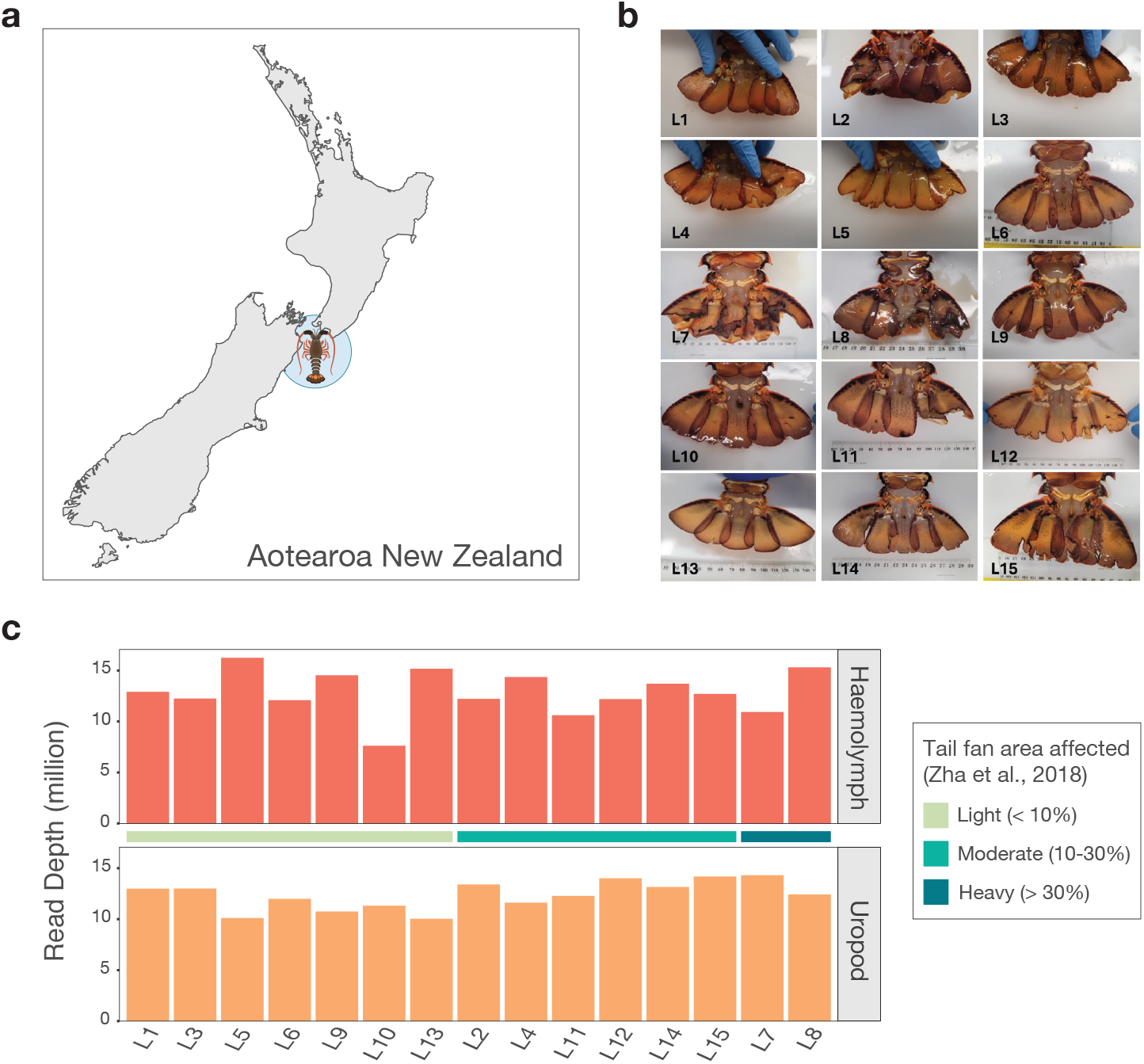
Sampling location, lobster tail fans, and RNA sequencing library depths. Map of New Zealand showing the area off the east coast of the South Island where lobsters were collected. B) Tail fans of lobsters 1 to 15 at the end of the holding period. C) Bar graphs showing the read depth of the 15 haemolymph samples (top) and uropod samples (bottom) in millions. Samples are ordered by the degree to which they were affected by TFN based on Zha et al. (2018b)’s categorisation: light green (Light; <10% tail fan area affected); green-blue (Moderate; 10 – 30%); and dark blue (Heavy; >30%).

### Assessment and scoring of tail fan damage

Lobsters were individually photographed for identification purposes and to keep track of tail fan damage progression from arrival until final assessment at the time of sampling (Figure 1b). Images were used to classify each individual into four different categories at the end of the holding trial based on the degree to which lobsters were affected by TFN including none: 0%, light: < 10%; moderate: 10 – 30%; and heavy: > 30% as per Zha et al. (2018b). The images together with the results from histopathology asssesment at the end of the holding period were compared against clinical and histology criteria presented in the case definition as per Jones et al. (2024).

Tail fan image analysis was carried out using the free software Image-J (Broeke et al., 2019), which allowed the identification and measurement of the area of the tail fan affected by blisters, melanisation, and tissue loss. The area for each of these conditions was expressed as a percentage of the total tail fan area. Since these were the main clinical signs observed in the tail fan, the sum of their area gives an estimation of the total area affected. Additionally, a tail fan impact score (TF-IS) was calculated, which considered different weighting for the three types of damage based on their impact on the lobsters’ tail fan (i.e., 50% to tissue loss area – A_tl_, 33% to melanisation area – A_m_, and 17% to blisters area – A_b_) as follows: TF-IS: (A_tl_ × 50 + A_m_ × 33 + A_b_ × 17)/100.

### Uropod and haemolymph sample collection

On the day of sampling, lobsters were individually sedated via exposure to cold temperatures inside a box with bagged ice for up to two hours. Haemolymph samples were taken from the base of the fifth walking leg using a pre-chilled 3mL syringe and a 25 G 1 ½ (0.5, 38mm) BD PrecisionGlideTM needle. Lobsters were then euthanised by pithing using a sterile knife. Small sections (0.5mm^2^) of affected uropods were cut using sterile scissors. Haemolymph and uropod tissue were stored separately in 2mL cryotubes containing 1 mL of RNA*later* and stored in a −80 °C freezer.

### RNA extraction and sequencing

Frozen haemolymph and uropod samples from 15 individual lobsters that exhibited clinical signs of TFN were defrosted and placed in ZR BashingBead Lysis Tubes (0.1 and 0.5 mm) (Zymo Research) filled with 1 mL of fresh DNA/RNA shield (Zymo Research) using sterile forceps. Lysis tubes were homogenised for five minutes in a mini-beadbeater 24 disruptor (Biospec Products Inc.) in one-minute intervals followed by one minute on ice. Total RNA was extracted from the 30 samples following the ZymoBIOMICS MagBead RNA kit (Zymo Research). RNA concentrations were quantified using a NanoDrop Spectrophotometer (ThermoFisher). The Illumina Stranded Total RNA Prep with Ribo-Zero plus kit (Illumina) was used for library preparation. Libraries were sequenced on the Illumina NovaSeqX platform and 150 bp paired-end reads were generated.

### Lobster metatranscriptome assembly and annotation

Trimmomatic (v0.38) (Bolger et al., 2014) was used to trim the Nextera paired-end adapters from the raw sequencing data. Using a sliding window approach, bases below a quality of five were trimmed with a window size of four. Bases were cut if below a quality of three at the beginning and end of the reads. Read library quality was assessed with FastQC to ensure the removal of adapters (https://www.bioinformatics.babraham.ac.uk/projects/fastqc/). Trimmed reads were then assembled *de novo* using MEGAHIT (v1.2.9) (Li et al., 2015). A sequence similarity-based approach was used to annotate assembled contigs against NCBI’s nucleotide (nt) database using the BLASTn algorithm (Camacho et al., 2009) and non-redundant (nr) protein database using DIAMOND BLASTx (v2.02.2) (Buchfink et al., 2021) with the “more-sensitive” flag set, and both ran with e-value cutoffs of 1×10^−5^.

### Discovery and abundance estimation of a novel virus in the *Nyamiviridae*

Viral hits were screened manually to identify invertebrate host-infecting viruses in uropod and haemolymph metatranscriptomes. Contigs annotated as viral (“Virus”) in the BLAST outputs with e-values <1×10^−10^ were inspected with additional BLASTn and BLASTp searches using the online BLAST tool to eliminate false positive hits. Putative viruses were considered if their top BLASTp hits were to previously known invertebrate-infecting viruses, particularly those infecting crustacean hosts. Genomes of putative viruses were recovered based on genome segments and lengths of related viruses.

The abundance of a novel crustavirus, Red rock lobster crustavirus (RRLCV) (family: *Nyamiviridae*), was estimated by mapping sequencing reads from each haemolymph and uropod library to the virus genome using Bowtie2 (Langmead & Salzberg, 2012). Read counts were standardised to reads per million (RPM) by multiplying library-standardised read counts by one million. Differences in crustavirus abundances between the three TFN categories proposed by Zha et al. (2018b) were tested using a Kruskal-Wallis test in R. Potential relationships between increasing tail fan damage and viral abundance were assessed using linear models and Spearman correlation (see GitHub).

### Data-mining for additional crustavirus sequences

To add context to the *Crustavirus* genus, the full RNA-dependent RNA polymerase (RdRp) sequence from the novel was used as bait to mine the NCBI Transcriptome Shotgun Assembly (TSA) database. All invertebrate metagenome (taxid: 1711999) and arthropods (taxid: 6656) transcriptomes were screened using the online translated Basic Local Alignment nucleotide search tool (tBLASTn) (https://blast.ncbi.nlm.nih.gov/Blast.cgi). The BLOSUM45 matrix was used to increase the chance of finding divergent viruses. Any putative viral polymerase hits were downloaded, translated, and subject to further query using BLASTp searches for confirmation.

### *Nyamiviridae* phylogenetic analysis

To place the novel lobster virus into its evolutionary context, a maximum-likelihood tree of the *Nyamiviridae* family was generated using IQ-TREE (v1.6.12) (Nguyen et al., 2015). Representative RdRp sequences from each genus in the *Nyamiviridae* were collected from NCBI’s Taxonomy database (https://www.ncbi.nlm.nih.gov/taxonomy), amino acid translations were combined with the additional TSA-mined *Crustavirus* species RdRps and aligned using the L-INS-i algorithm in MAFFT (v7.450) (Katoh & Standley, 2013). Alignments were manually inspected to observe the alignment of conserved motifs and ambiguously aligned regions were trimmed using trimAl (v1.2) (Capella-Gutiérrez et al., 2009) with the “automated1” flag set. IQ-TREE was used to generate a phylogeny based on this alignment with the ModelFinder “MFP” flag set to select the best-fit model of amino acid substitution and 1000 ultra-fast bootstrapping replicates (Hoang et al., 2018). The “alrt” flag was also added to perform 1000 bootstrap replicates for the SH-like approximate likelihood ratio test (Guindon et al., 2010). The resulting phylogenetic tree was rooted at its midpoint and annotated in FigTree (v1.4.4) (http://tree.bio.ed.ac.uk/software/figtree/).

### PCR confirmation of RRLCV in lobster tissues

Primers were designed to confirm amplicons from extracted haemolymph and uropod RNA using reverse transcription polymerase chain reaction (RT-PCR). Primers were designed in Geneious Prime (v2020.2.4) using the L protein as the target. In the Design New Primer function, three primer sets (forward and reverse) were designed for the PCR with the following attributes: 25 bp in length, GC contents 40-60%, and T_m_ of 51-52°C. The resulting primer sets generated products 160-187 bp in length, covering regions across the polymerase with no overlap or repetition (Supplementary Data 2 Table 1). An additional three reverse primers were designed for the initial cDNA step, complementary to the negative-sense viral genome and overlapping the reverse primers of the PCR step in order to selectively increase the quantity of RRLCV viral cDNA generated for the PCR.

**Table 1.** Characterisation of the tail fan damage for the 15 lobsters sampled at the end of the holding trial. Descriptive categories (Light, Moderate, and Heavy) are presented and the corresponding percentage (%) of the tail fan affected by blisters, melanisation, and tissue loss is also reported. A final tail fan impact score was calculated for each lobster that considers the relative impact of blisters, melanisation, and tissue loss.

| **Lobster** | **Zha et al. 2018b category** | **Blisters (%)** | **Melanisation (%)** | **Tissue loss (%)** | **Total area compromised (%)** | **Score** |
| --- | --- | --- | --- | --- | --- | --- |
| 1 | Light | 0.00 | 0.00 | 0.75 | 0.75 | 0.38 |
| 2 | Moderate | 8.66 | 5.12 | 12.66 | 26.44 | 9.49 |
| 3 | Light | 0.00 | 2.46 | 3.77 | 6.24 | 2.70 |
| 4 | Moderate | 6.97 | 3.86 | 8.68 | 19.50 | 6.80 |
| 5 | Light | 0.79 | 3.35 | 5.43 | 9.57 | 3.95 |
| 6 | Light | 0.00 | 0.76 | 1.24 | 2.00 | 0.87 |
| 7 | Heavy | 2.95 | 25.28 | 25.13 | 53.35 | 21.41 |
| 8 | Heavy | 5.47 | 15.82 | 17.04 | 38.33 | 14.67 |
| 9 | Light | 0.00 | 1.13 | 4.68 | 5.81 | 2.71 |
| 10 | Light | 1.64 | 0.34 | 2.71 | 4.69 | 1.74 |
| 11 | Moderate | 2.17 | 4.17 | 17.38 | 23.72 | 10.43 |
| 12 | Moderate | 0.00 | 2.58 | 6.87 | 9.45 | 4.29 |
| 13 | Light | 0.00 | 0.00 | 0.75 | 0.75 | 0.38 |
| 14 | Moderate | 5.58 | 1.50 | 6.42 | 13.50 | 4.65 |
| 15 | Moderate | 2.91 | 3.15 | 7.34 | 13.40 | 5.20 |

RNA was converted to cDNA using a High-Capacity cDNA Reverse Transcriptase kit (Applied Biosystems, ThermoFisher Scientific) with the modification of 1 μL of an equal mixture of the three cDNA reverse primers and 1μL of random hexamer primers per reaction (2 μL of primers total). PCR amplification was performed with AllTaq Master Mix (Qiagen) and a thermocycler protocol as follows: 95 °C for 2 min; 95 °C for 5s; 40 cycles of 94 °C for 5 s, 50 °C for 15 s, 72 °C for 30 s; 72 °C for 10 min. Gel electrophoresis with fluorophore (SYBR Safe DNA Gel Stain, Invitrogen) in 1% agarose (Invitrogen) at 70V for 65 min was used to separate PCR products, including negative controls and a 100 bp ladder (Invitrogen), on 0.5X TAE running buffer.

PCR bands were visualised by UV light using GelDoc Go Imaging System (Bio-Rad Laboratories). Bands of expected sizes for each of the three primer sets were excised, and DNA was purified using the Qiagen QIAquick PCR and Gel Cleanup Kit with 30 μL of EB buffer used at the end to elute DNA from the column. DNA was sent for bi-directional Sanger sequencing at the University of Otago Genetic Analysis Services, New Zealand. Sequences were aligned against the RRLCV L protein in Geneious Prime (v2023) to confirm that expected sequences and lengths were obtained. Primers and Sanger sequencing were also screened against the online NCBI nt database to rule out non-specific amplification or contamination.

### Cellular microbe characterisation and analysis

Cellular microbes (prokaryotes, and eukaryotes) present in each library were classified using CCMetagen (Marcelino et al., 2020). Reads were first mapped to the NCBI nucleotide database, excluding unclassified environmental microbes and cloning vectors using k-mer alignment (KMA) (Clausen et al., 2018), ensuring a match was only counted if both paired-end reads were mapped to a reference. KMA outputs were then processed with CCMetagen to produce microbial taxonomic classifications of uropod and haemolymph samples and combined at the genus level. The composition (presence and abundance) of lobster uropod and haemolymph microbiomes in the context of damage caused by TFN was explored. All analysis was performed using R (v4.11). CCMetagen outputs were imported into R and filtered to exclude non-microbial eukaryotes and miscellaneous viruses, as well as genera that were present in less than three samples.

Alpha diversity of the microbial population was considered based on genus richness (number of genera per sample) and Shannon diversity, which accounts for both richness and evenness (abundance). Statistically significant differences in alpha diversity between TFN categories were tested using analysis of variance (ANOVA). Relationships between measurements/scoring of tail fan damage and these alpha diversity measures were also determined using linear models.

Beta diversity of lobster microbiomes based on tail fan damage and TFN categorisation was assessed using non-metric multidimensional scaling (NMDS). A distance matrix was generated using the vdist function available in the vegan package (Oksanen et al., 2022) with Bray-Curtis dissimilarity as the distance measure. NMDS was performed on the distance matrix using the metaMDS function from vegan and permutational multivariate analysis of variance (PERMANOVA) with the adonis2 function was used to test for statistical significance of the effect of TFN category on microbiome similarities. Results were graphed using ggplot2 (Wickham, 2016).

## Results

### Classification of TFN in spiny lobsters and sequencing overview

Lobsters received from commercial fisheries were suspected to exhibit TFN at the point of entry to the laboratory (i.e., erosion of the edges of the telson and/or uropods, some with the presence of blisters on the ventral side of telson and/or uropods). Based on the scale provided by Zha et al (2018b), on reception, the 15 lobsters were classified as follows: n=1 with no clinical signs of disease, and n=14 light TFN.

At the end of the holding period, histopathology assessment confirmed that all but three lobsters (L1, L6, and L13) met both, clinical and histology criteria presented by Jones et al (2024). Several lobsters suffered the loss of tissue from affected telson and/or uropod, with clear blackening (melanisation) usually in areas associated with preexisting blisters (Figure 1b). Based on the scale provided by Zha et al. (2018b), at the end of the holding period, the lobsters were classified as follows: n=7 of the lobsters sampled exhibited light TFN, n=6 moderate TFN, and n=2 heavy TFN (Table 1). Symptoms progressed from the base of the uropod and telson, with the tail fan area covered by blisters, melanisation, and tissue loss varying among individuals (Table 1). Blister area did not go over 9%, while melanisation was almost half of that for most individuals, except for the two most affected lobsters which had 25.3 and 15.8% of tail fan area covered, respectively (Table 1). Tail fan tissue loss also varied among individuals ranging from 0.8% to 25.1% (Table 1). The tail fan impact score ranged from 0.4 in the least affected individual to 21.4 in the most compromised individual.

RNA sequencing of haemolymph and uropod samples yielded 7.7 – 16.2 million reads per library (average: 12.6 million) (Figure 1) and an average of just under 350,000 contigs per assembled metatranscriptome (Supplementary Table S1).

### Identification and PCR confirmation of a novel crustavirus species

Contigs annotated as viral in the assembled haemolymph and uropod metatranscriptomes were screened for putative crustacean-infecting viruses present in multiple affected lobsters for their possible implication in TFN. From this, we identified a single likely lobster-infecting virus across all the uropod samples and 20% (3/15) of the haemolymph samples (Figures 2 and 3). We phylogenetically analysed the virus, putatively named Red rock lobster crustavirus (RRLCV), which fell into the *Crustavirus* genus within the *Nyamiviridae* family of unsegmented, negative-sense single-stranded RNA viruses (Figure 2). Viruses in this family were first identified in ticks (Arthropoda) and associated bird species, but their host range has since expanded to various other invertebrate host species (Kuhn et al., 2013). RRLCV was most closely related to another novel virus from a European green crab (*Carcinus maenas*) epidermis transcriptome (GFYW01) identified by screening the TSA database. The two viruses shared 67% pairwise identity. RRLCV also shared 61% amino acid identity with the previously known *Wenzhou Crab Virus 1* (YP_009304558.1), identified in several crab and barnacle species in China (Li et al., 2015) (Table 1). No other invertebrate-associated viruses of interest were identified across multiple lobsters.

**Figure 2.**
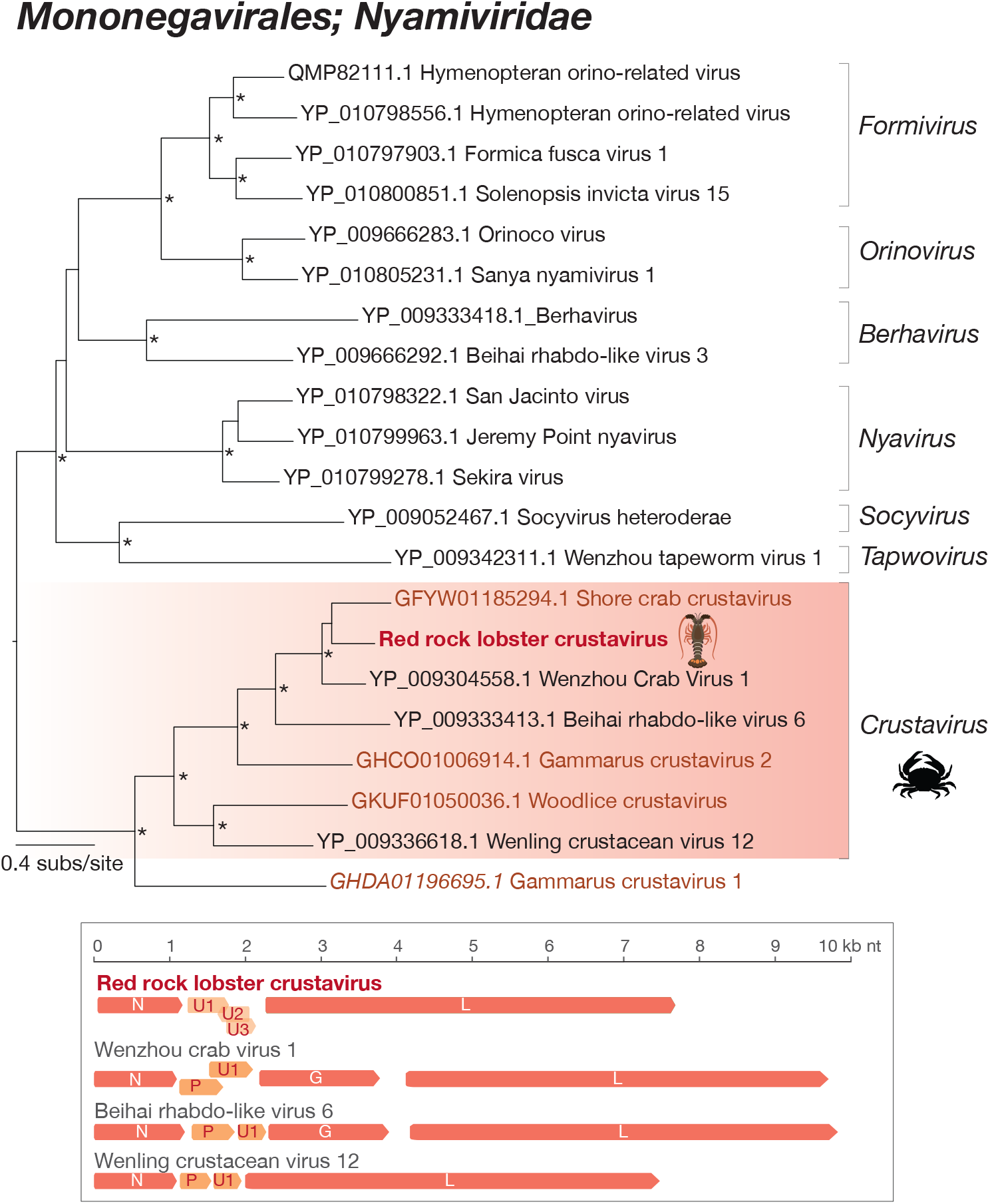
Maximum likelihood phylogenetic tree of RdRps from viruses in the *Nyamiviridae* and representative crustavirus genomes. The *Crustavirus* genus is highlighted in a red box and the novel RRLCV is indicated in bold red and by a lobster silhouette. Additional viruses identified through the mining of the TSA are indicated in brown and denoted by their host’s scientific name or general host. Amino acid substitutions per site are indicated by the key on the left-hand side of the tree. The tree is rooted at the midpoint and nodes with ≥ 95 UFbootstrap support values are denoted by an asterisk (*). Genome organisations of recognised crustaviruses and RRLCV are shown below.

**Figure 3.**
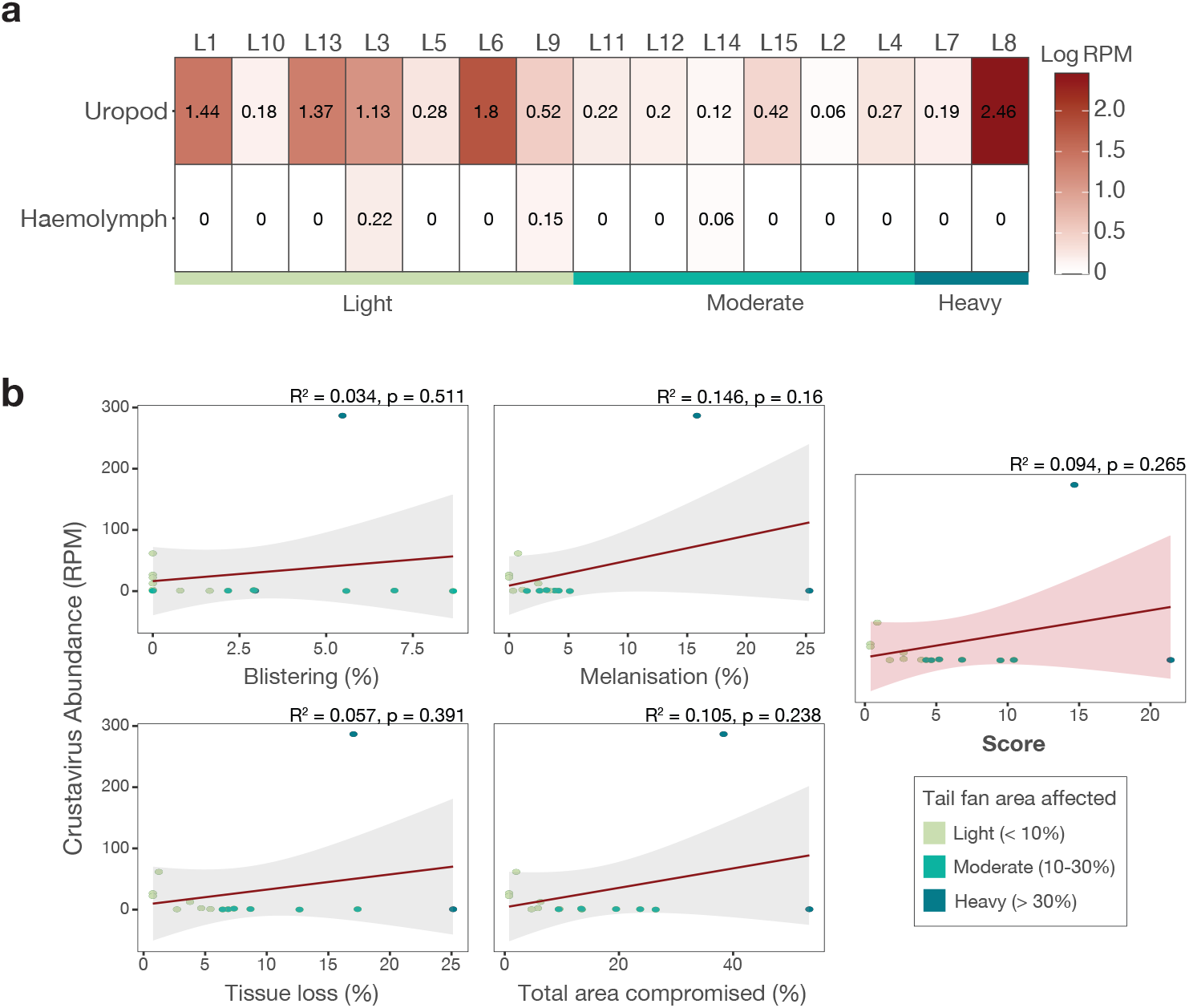
Abundance of RRLCV in lobster samples and correlation with tail fan damage. A) Heatmap of crustavirus abundance (logged RPM) in uropod and haemolymph samples, organised by Zha et al. (2018b)’s TFN categorisation. B) Correlation of crustavirus abundance (RPM) with the percentage of tail fan affected by blistering, melanisation, tissue loss, the total area compromised, and overall scoring of tail fan damage. None of the measurements or scoring of tail fan damage in the affected lobsters were significantly correlated with viral abundance.

We recovered a likely full 7,681 nucleotide genome of RRLCV (Figure 2). The genome included a 5,412 nucleotide L protein (RdRp), a 1,125-nucleotide nucleoprotein, and three predicted open reading frames encoding hypothetical proteins (unknown 1 to 3) ranging from 393 to 549 nucleotides. None of the hypothetical proteins shared significant sequence similarity with any known sequences in the NCBI nt and nr databases. Notably, we could not detect a gene encoding a glycoprotein. However, other viruses in this genus also lack this gene. Mining invertebrate transcriptomes in the TSA database using the L protein of RRLCV as bait revealed four additional invertebrate viruses: three from crustaceans – *Carcinus maenas* (GFYW01), *Gammarus pullex* (GHCO01), and *Gammarus fossarum* (GHDA01) – and one from an isopod host, *Trichoniscus matulici* (GKUF01) (Figure 2 and Table 2). All four viruses clustered with previously identified crustaviruses.

**Table 2.** Novel crustaviruses identified in this study.

| Virus | Host | GenBank accession | Blastx hit | Identity (%) | Complete? | Method |
| --- | --- | --- | --- | --- | --- | --- |
| Red rock lobster crustavirus | *Jasus edwardsii* | PQ440166 | YP_009304558.1 | 60.8 | Yes | RNA-seq |
| Shore crab crustavirus | *Carcinus maenas* | GFYW01 | YP_009304558.1 | 65.8 | No | TSA mining |
| Gammarus crustavirus 1 | *Gammarus fossarum* | GHDA01 | YP_009336618.1 | 33.8 | No | TSA mining |
| Gammarus crustavirus 2 | *Gammarus pulex* | GHCO01 | YP_009304558.1 | 39.6 | No | TSA mining |
| Woodlice crustavirus | *Trichoniscus matulici* | GKUF01 | YP_009336618.1 | 40.4 | No | TSA mining |

RRLCV was most abundant in a lobster with heavy TFN (at 287 RPM) and four other lobsters with light TFN (Figure 3a). The virus was found at low abundances in all other uropod samples and only three haemolymph samples. Viral abundance did not differ significantly between TFN severity groups as proposed by Zha et al. (2018b) in the uropod (Supplementary Figure S1a, Kruskal-Wallis χ^2^ p-value 0.104). Similarly, there were no significant relationships between increased viral abundance and increased blistering, melanisation, tissue loss, or total area compromised (as percentages of the full uropod) (Figure 3b). The overall scoring of tail fan damage was not significantly correlated with viral abundance (Figure 3b). We confirmed the presence of RRLCV in uropod and haemolymph samples by PCR, with all 15 lobsters testing positive in both sample types (Supplementary Data 2).

### Lobster cellular microbiomes

We found lobster tissue microbiomes to be comprised of 30 cellular (non-viral) microbial genera (Supplementary Figure S2) falling within 15 taxonomic orders (Figure 4). Lobsters carried between zero to 15 different microbial orders. Flavobacteriales was the most widespread and abundant order, being found in all uropod samples, followed by Saprospirales in 60% of uropod samples. Microbial groups of interest due to their past association with TFN, such as *Vibrio* sp. and *Aquimarina* sp., were identified in three to six (20 to 40%) of the affected lobsters. Lobsters in the moderate to heavy group carried 5 to 6.6 microbial orders on average while the light TFN group carried an average of 4.4. Only two orders were detected in haemolymph samples, Micrococcales (in 73% of samples) and Chroococcidiopsidales (in 20%).

**Figure 4.**
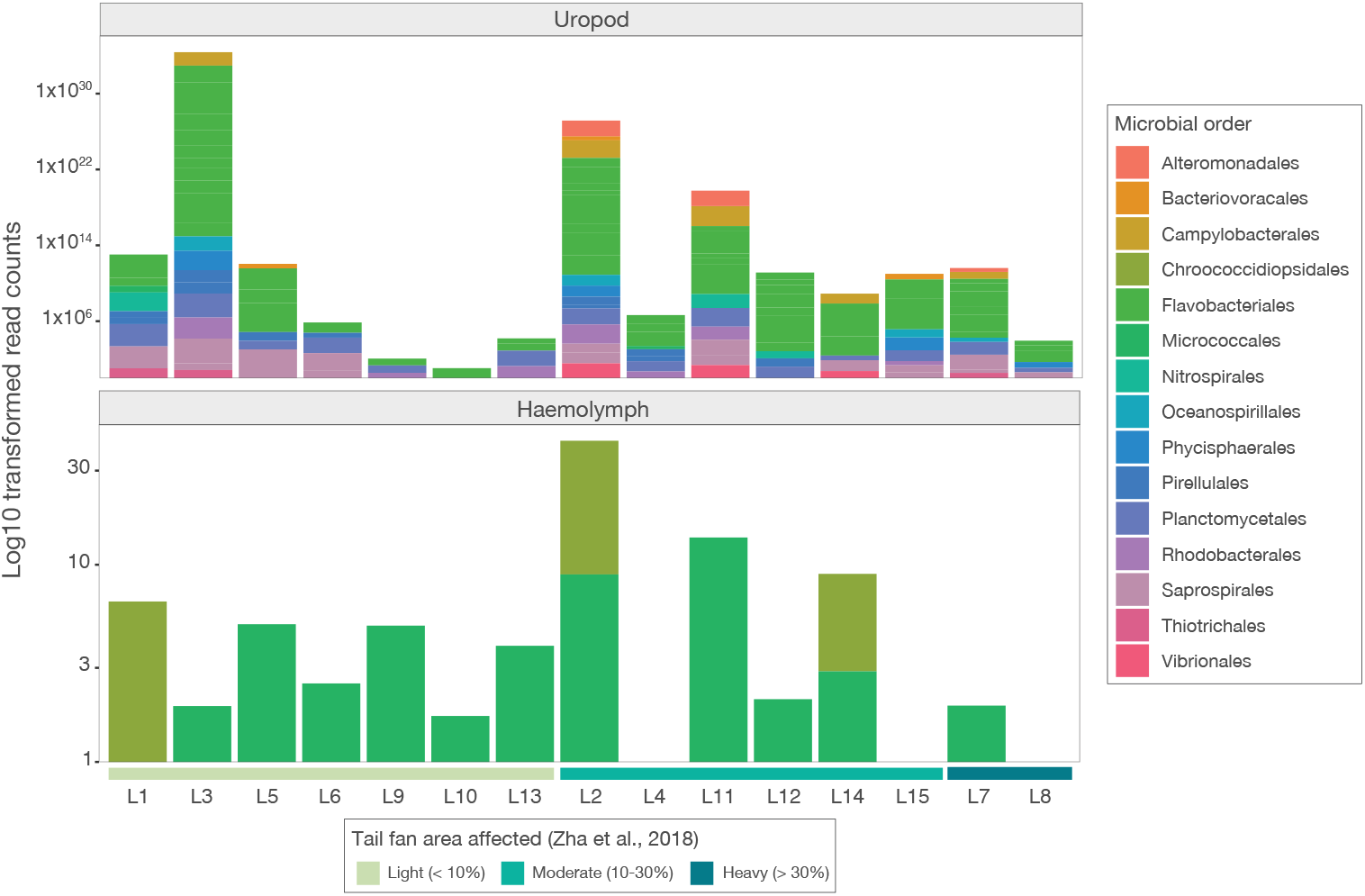
Bar plots showing abundances of 15 microbial orders found across the samples. Uropod (top) and haemolymph (bottom) samples are grouped by the degree to which they were affected by TFN, and abundance counts are log10 transformed.

Overall, genus richness and Shannon diversity of lobster microbiomes were significantly different between sample types (uropod and haemolymph) (Figure 5a, ANOVA p-values 2.1×10^−6^ and 0.009, respectively). Shannon diversity differed significantly between TFN- affected groups when controlling for sample type (*p* = 0.033), where moderately and heavily affected lobsters had higher genus-level Shannon diversity than lobsters with light TFN. Microbial communities also differed significantly between sample types (Figure 5b, PERMANOVA p-value 0.001). Nevertheless, microbiome compositions were not significantly different between TFN severity groups in either tissue type (*p* = 0.348). We also evaluated possible correlations between alpha diversity measures and the percentage of the total tail fan affected by the four factors (blistering, melanisation, tissue loss, and total area compromised) and found no significant relationship for any of the factors (Supplementary Figure S3).

**Figure 5.**
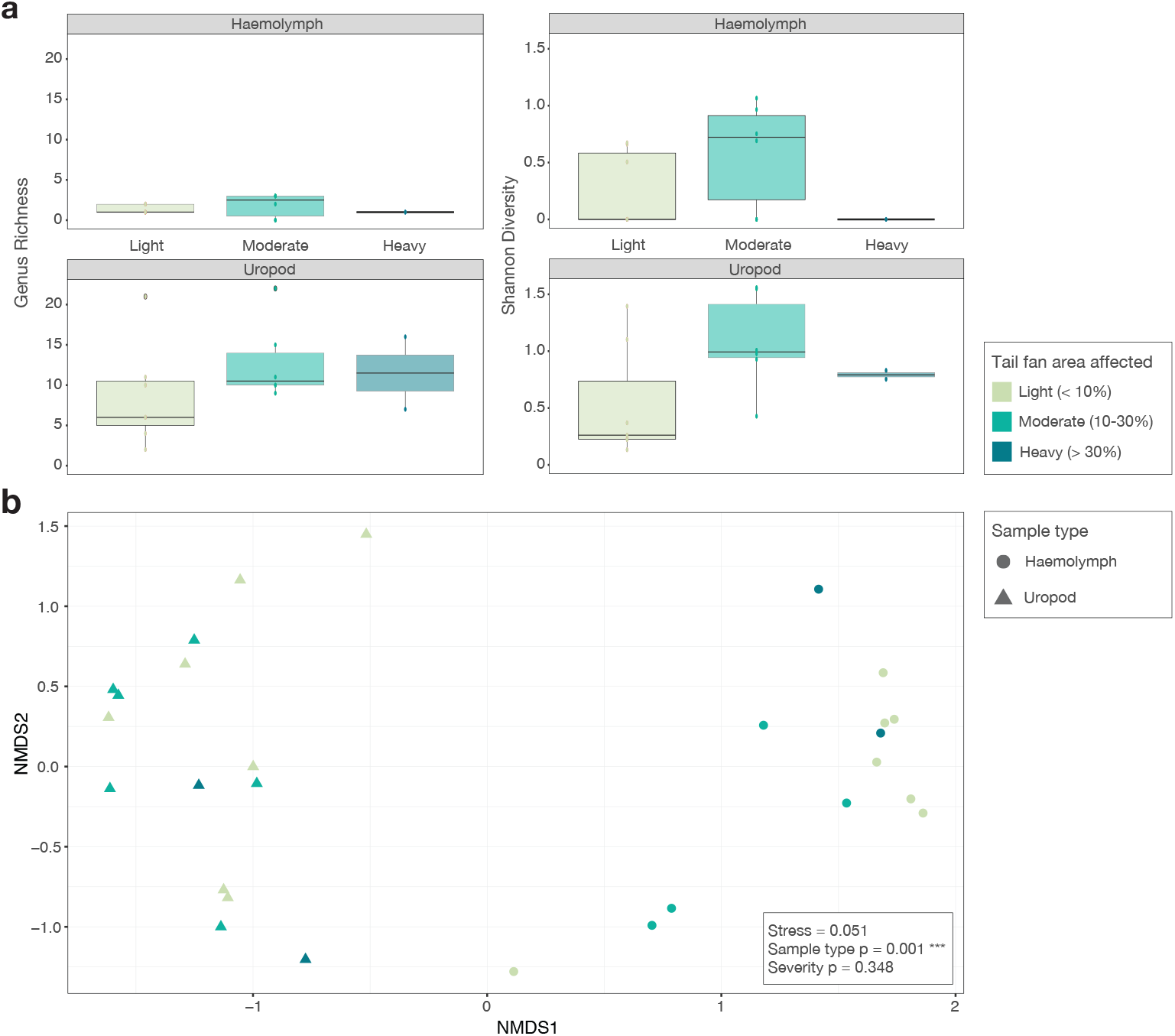
Alpha and beta diversity of lobster microbiomes. A) Genus richness of haemolymph and uropod samples by area affected (left) and Shannon diversity of haemolymph and uropod samples by area affected (right). Richness (genus-level) differed significantly between sample types, but not by the degree to which the tail fans were affected (ANOVA tail fan area affected group p-value: 0.368; sample type p-value: 2.07×10^−6^) and Shannon diversity differed significantly by TFN group and sample type (ANOVA tail fan area affected group p-value: 0.033; sample type p-value: 0.009). B) Non-metric multidimensional scaling plot based on Bray-Curtis dissimilarity of haemolymph and uropod microbiomes. Sample type significantly influenced microbiome composition (PERMANOVA p-value = 0.001) while the area affected by TFN did not (PERMANOVA p-value = 0.348).

## Discussion

To investigate potential disease-causing microbial candidates associated with TFN, we have used metatranscriptomics to describe the microbial communities associated with uropod and haemolymph samples from affected lobsters. We report the presence of a new crustavirus species within the uropod of all sampled red rock lobsters with TFN (n=15), named Red rock lobster crustavirus (RRLCV). We identified four additional novel viruses in the same genus by mining published Arthropoda transcriptomes. Crustaviruses (family: *Nyamiviridae*) infect crustaceans but have not been previously associated with disease (Dietzgen et al., 2021).

The closest relative of RRLCV was identified in an epidermis transcriptome from a European green crab (*C. maenas*) with no noted signs of disease (Oliphant et al., 2018), suggesting these viruses infect tissues that make up the carapace and tail fans of crustaceans and can do so asymptomatically. *Carcinus* species are also susceptible to unique shell diseases, including those causing ulceration and melanisation, linked to the enzymatic activity of chitinolytic bacteria (Mancuso, 2013). Only one other virus has been associated with TFN in spiny lobster in New Zealand: *Armadillidium vulgare* iridescent virus (family: *Iridoviridae*) (Pande et al., 2021), although it is unclear what the genetic relatedness of the iridoviral sequences in those studied lobsters was to the original virus, which normally infects the isopod *Armadillidium vulgare*, causing infected animals to turn iridescent (Federici, 1980). Irido-like DNA viruses have been found in other crustaceans and have been linked to mortalities in amphipods and European shore crabs (Subramaniam et al., 2020); however, we did not find any DNA viral transcripts in the lobsters sampled here and previously identified viruses in crustaceans and isopods do not cause similar shell diseases.

Our analysis did not reveal a significant clustering of lobsters into groups (light, moderate, heavy) based on viral abundance, nor did we find a significant positive relationship between measures of TFN damage and viral abundance. RRLCV was only detected reliably, and at relatively high abundances, in uropod samples in the RNA-seq data, which would be consistent with a virus able to cause pathology in the tail fan. We detected viral sequences in both uropod and haemolymph samples on PCR. Consistent detection of RRLCV polymerase sequences in haemolymph in PCR but not RNA-seq suggests the presence, but not active or efficient replication, of RRLCV sequences in haemolymph. Haemolymph and the blisters, unique to TFN in lobsters from Oceania, are both assumed to be sterile, or free from culturable bacteria, implying TFN arises from external infection (Diggles et al., 1999; Jones et al., 2024; Musgrove et al., 2005). The inconsistent detection of RRLCV in the haemolymph in RNA-seq data may further suggest the virus is acquired externally and does not become systematic, at least in the early stages of an infection. Still, the route of infection and tissue tropism of crustaviruses are currently unknown and limit our ability to infer patterns of viral abundance across tissues and disease progression.

Several factors may influence the estimated abundance of a virus in a sample (Louten, 2016) and therefore the ability to correlate this with disease “severity”. At the time of sample collection and RNA extraction, we did not know what microbial agents might be identified, nor the proportion of potentially infected tissue type(s) in each sample. For example, variations in the ratio of the epidermis (where the virus may reside) to muscle or nervous tissue could significantly affect the estimated viral load, leading to inconsistencies that obscure potential relationships between the degree of tail fan damage and viral load. Variability in the stage of infection across individuals is another possible confounding factor. In some lobsters, the virus may have been in the early stages of infection, localised in the uropod, while in others, it may have been progressing or resolving. Environmental factors and stressors have also been implicated in TFN. Water quality, temperature, and handling (Freeman & MacDiarmid, 2009; Musgrove et al., 2005; Zha et al., 2018a, 2019) as well as host factors, such as genetics, age, and prior health status could modulate a lobster’s immune response and therefore viral replication. Finally, the scoring system based on photo analysis and the categorisation system by Zha et al. (2018b) both rely on visual assessments that may not fully capture the biological complexity of TFN or viral infection. These metrics would benefit from integrating them with factors such as molecular markers of immune response, histopathological analyses, and more deliberate and careful sampling of tissue type and stage of infection to provide a more accurate picture of TFN progression and its possible association with viral infection.

We were unable to detect a viral glycoprotein in RRLCV. Glycoproteins are used by many viruses for host cell attachment and fusion and to evade host immune responses (Banerjee & Mukhopadhyay, 2016). Viruses within the family *Nyamiviridae* generally encode a glycoprotein in their genome (Dietzgen et al., 2021). However, several other species within the *Nyamiviridae* lack glycoproteins, including the crustaviruses *Behai rhabdo-like virus 5* and *Wenling crustacean virus 12* (Shi et al., 2016), and a nyavirus identified in ticks (Kobayashi et al., 2021). Therefore, while a glycoprotein may not have been detected due to a lack of sequence similarity with other known crustavirus glycoproteins, it would not be unusual for a crustavirus not to encode one and suggests RRLCV may have an alternative mode of host cell invasion, such as lipid-membrane interactions or endocytosis.

Shell diseases in decapods have more frequently been associated with bacteria (Chistoserdov et al., 2012). Still, TFN in New Zealand lacks a consistently identified causative bacterial pathogen. We characterised the wider “infectomes” – infectious agents – of the lobsters to identify any suspect prokaryotic or eukaryotic microorganisms. Classical shell diseases have been linked to the chitinolytic, proteolytic, and lipolytic activity of some bacteria (Shields, 2011; Zha et al., 2018a). We found 30 bacterial genera, including *Aquimarina, Vibrio, Tenacibaculum*, and *Flavobacteria* in the uropod samples which have previously been associated with TFN (Chistoserdov et al., 2012; Zha et al., 2019). *Aquimarina* species have been noted as abundant marine microorganisms and potential first colonisers of the uropod after it becomes compromised (Meres et al., 2012) and have been associated with cuticular diseases in cultured larvae (Ooi et al., 2020). *Vibrio* species are also frequently identified (Porter et al., 2001). *Vibrio* and *Aquimarina* species are thought to cause shell disease symptoms in American and spiny lobsters (Chistoserdov et al., 2012; Geddes et al., 2004), while others link these diseases to the presence of multiple bacteria (Zha et al., 2019). Other implicated genera, like *Shewanella, Photobacterium, Pseudomonas*, and eukaryotic pathogens, such as *Enterospora* sp. (Musgrove et al., 2005; Pande et al., 2021; Zha et al., 2018a), were not identified in the metatranscriptomes of the lobsters here, but the presence of *Photobacterium* has been confirmed in media cultures from uropod swabs of two lobsters before sampling. Like these other studies, we did not find any single bacterial genus or species consistently across all affected lobsters and most are generally associated with the normal flora of otherwise healthy lobsters (Porter et al., 2001) or are present to some degree regardless of TFN status (Meres et al., 2012). This makes it difficult to determine potential pathogenicity without further analysis or challenge studies. Haemolymph samples were mostly free of microbes. Crustaceans can rapidly clear microbes from their haemolymph (Martin et al., 1993) and it is considered sterile (Jones et al., 2024), therefore our findings are consistent with the general lack of culturable microbes obtained from lobster haemolymph.

It has been proposed that a general dysbiosis of the microbial flora associated with the lobsters may contribute to susceptibility to TFN (Rowley & Coates, 2013). For example, several studies have noted lower microbial diversity but higher microbial variability, as well as an increased abundance of chitinolytic, proteolytic, lipolytic, and melanin-producing strains of bacteria in affected samples compared to unaffected samples (Shields, 2011; Zha et al. 2018a, 2019). We found that heavily affected lobsters had higher microbial alpha diversity in their uropod compared to haemolymph but the lack of unaffected (control) lobsters in this sample set prevented comparison.

We provide the first report of a novel microbial agent consistently associated with the uropods of New Zealand red rock lobsters with TFN – a novel *Crustavirus* species. As we only examined affected lobsters, future work should expand to test symptomatic and asymptomatic individuals to determine if RRLCV associates closely with the disease and what its distribution and tissue tropism may be. TFN is a complex, likely multifactorial disease that requires more detailed investigation. While the tail fan has been the primary focus of much research into TFN, pathology and blistering have been detected on the carapace, walking legs, as well as in the gills and some internal organs (including the mid-gut and heart) of some lobsters (Diggles et al., 2022; Zha et al., 2018b). This points to a potentially more systematic issue beyond mere physical damage during handling and subsequent bacterial invasion. If RRLCV appears to be closely associated with the disease, further investigation of the ecology, transmissibility, and pathogenicity of this virus is warranted.

## Acknowledgements

We wish to thank the North Island Cray Management Area Council (CRAMAC) for sourcing the lobsters for this project. The authors are thankful to Joanna Copedo (histopathologist, Cawthron Institute) that assisted with histological assessment of the lobster’s tail fan. Thanks to Dr Stephanie Waller for her lobster illustration.

## Data availability

Raw sequence reads are available on the NCBI Short Read Archive (SRA) under the BioProject accession PRJNA1219689. The RRLCV genome sequence is available under the GenBank accessions PQ440166. Extended data and R scripts used to generate results and figures in this study can be found on GitHub: https://github.com/maybec49/Lobster_TFN.

