## Supplementary Figures for "A novel crustavirus associated with tail fan necrosis in New Zealand red rock lobsters, *Jasus edwardsii*"

### **Supplementary Material**

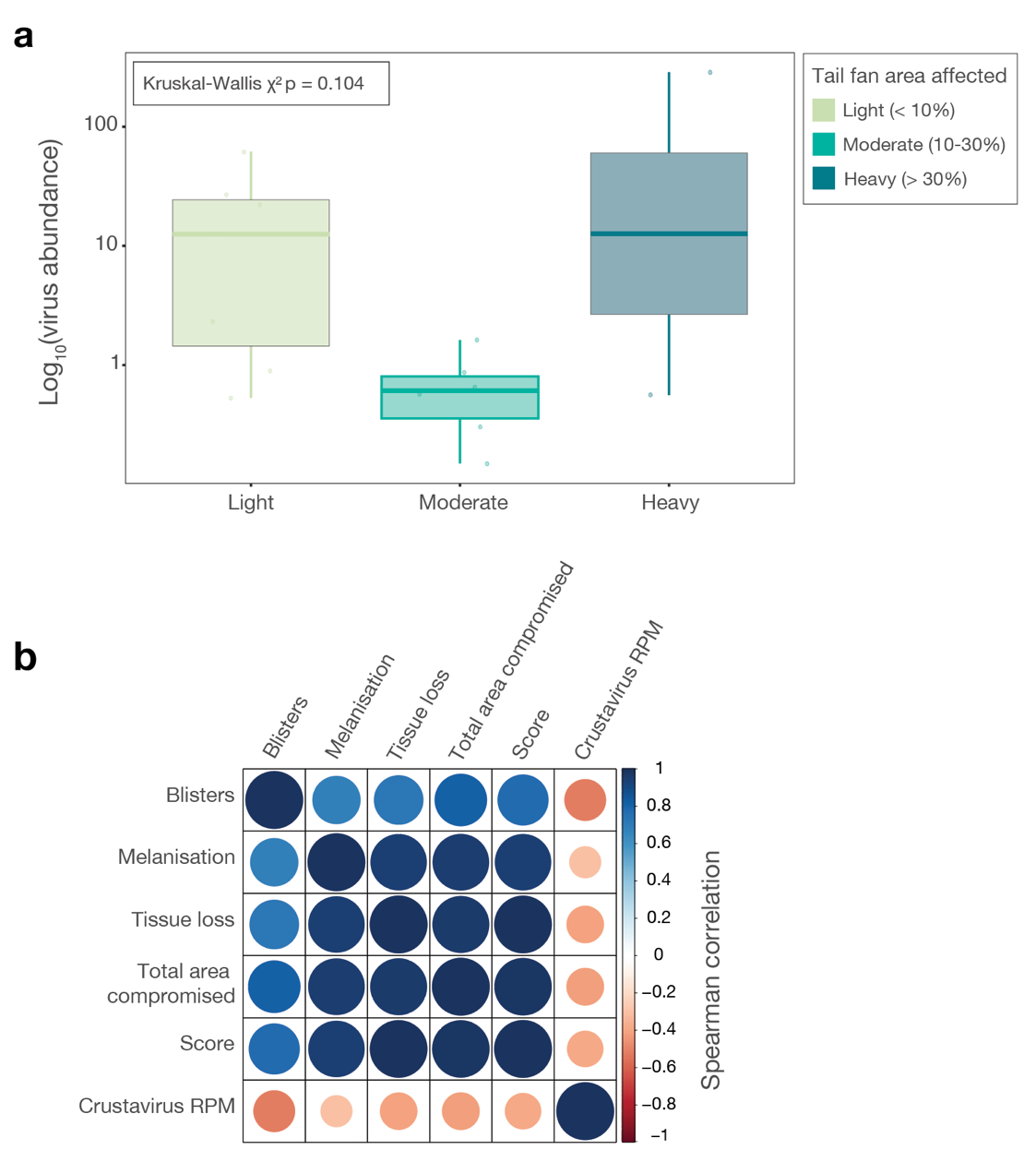

Figure S1. Association of viral abundance with categorisation of TFN and measures of tail fan damage.

A) Box and whisker plots of log10 crustavirus abundance in uropod samples by TFN. There was no significant difference in crustavirus abundance between affected groups as outlined by (Zha et al. 2018) in uropod samples (Kruskal-Wallis χ² p-value 0.104). B) Spearman correlation of measures of tail fan damage with crustavirus abundance (RPM). There were weak, but insignificant, correlations between viral abundance and blistered, melanisation, tissue loss, and total area compromised (as percentages of the total tail fan area), as well as with the overall damage scoring.

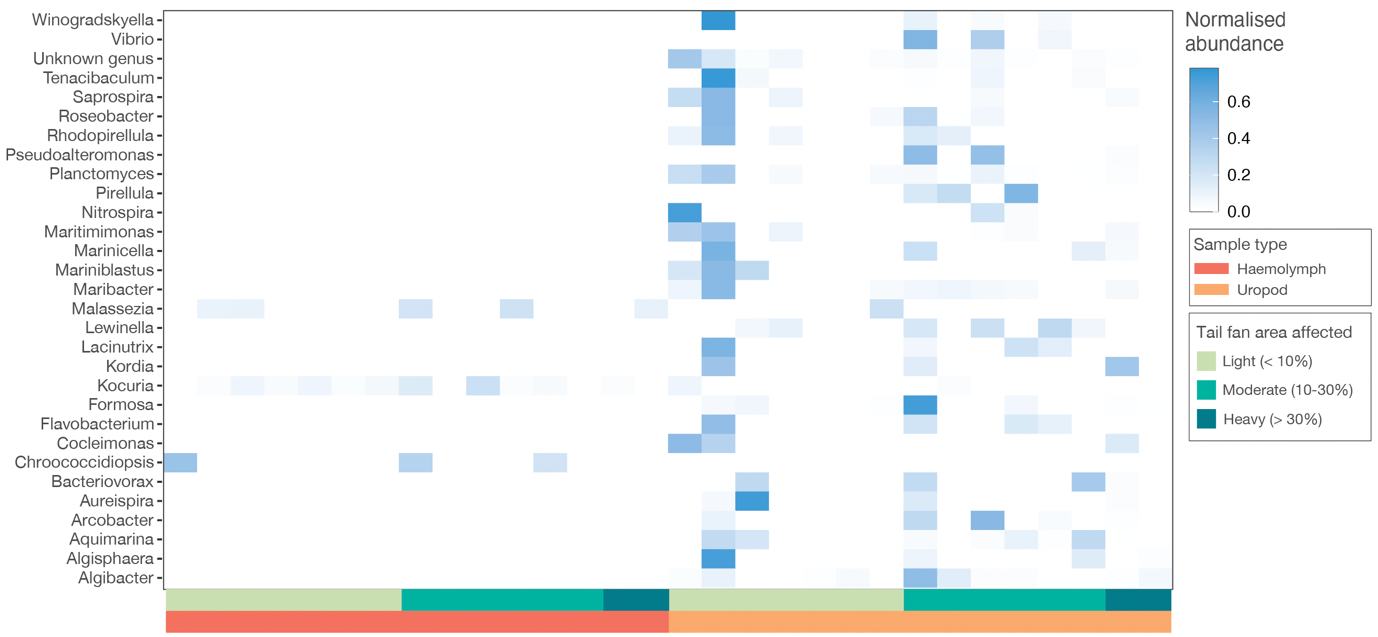

Figure S2. Heatmap showing abundances of the key microbial genera comprising the lobster microbiomes.

Samples are ordered by sample type (haemolymph – dark orange and uropod – orange) and by percentage of tail fan affected by TFN (light – green, moderate – light blue/green, and heavy – dark blue). Abundances are normalised by microbe genus.

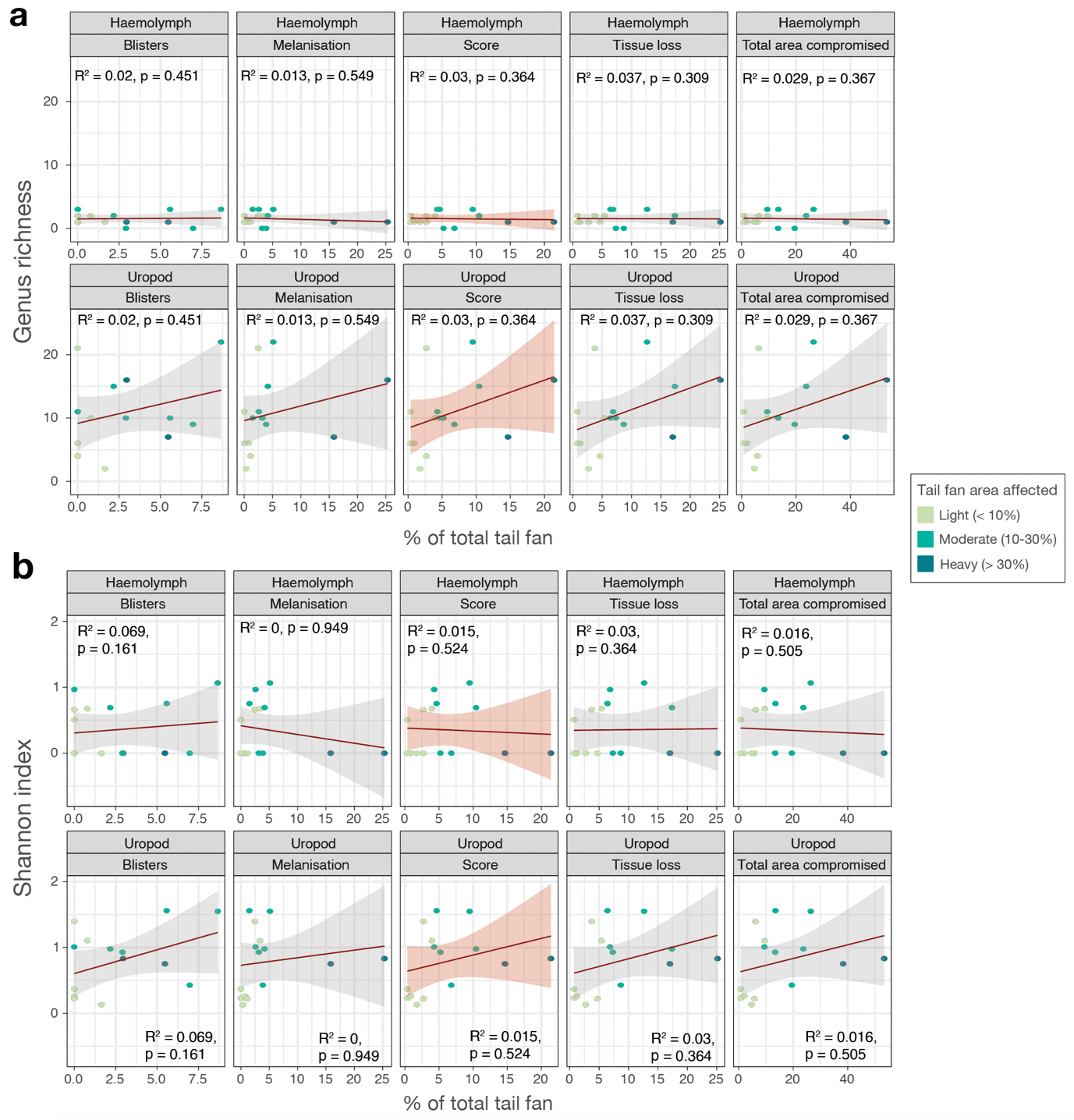

Figure S3. Relationships between microbial alpha diversity and percentage of tail fan damage.

A) Correlation of sample bacterial richness with percentage of tail fan affected by blistering, melanisation, tissue loss, and total area compromised and overall scoring of tail fan damage in haemolymph and uropod (top). B) Correlation of sample Shannon diversity with percentage of tail fan affected by: blistering, melanisation, tissue loss, and total area compromised and overall scoring of tail fan damage in haemolymph and uropod (bottom). None of the measurements or scoring of tail fan damage in the affected lobsters were significantly correlated with alpha diversity of the bacterial communities in haemolymph or uropod.
