## Supplementary Data 2 for "A novel crustavirus associated with tail fan necrosis in New Zealand red rock lobsters, *Jasus edwardsii*"

**Supplementary Table 1.** RRLCV PCR primers sequences and product lengths.

| **Primer set name** | **Forward seq** | **Reverse seq** | **Product length (bp)** |
| --- | --- | --- | --- |
| A | GTATAGATGAGTGTTGTGCATC | CTGGTTAGTAAGATGTCACCTT | 160 |
| V | TGGCCTAGAGAGTATGATATGA | GCCATCTGTGATTCTCTAGATT | 173 |
| T | TACATCATCCTCTGTCCTTAGA | AAAGTCTCCTCGTACTTTATCC | 187 |
| Complementary A | - | GACCAATCATTCTACAGTGGAA | NA |
| Complementary V | - | CGGTAGACACTAAGAGATCTAA | NA |
| Complementary T | - | TTTCAGAGGAGCATGAAATAGG | NA |

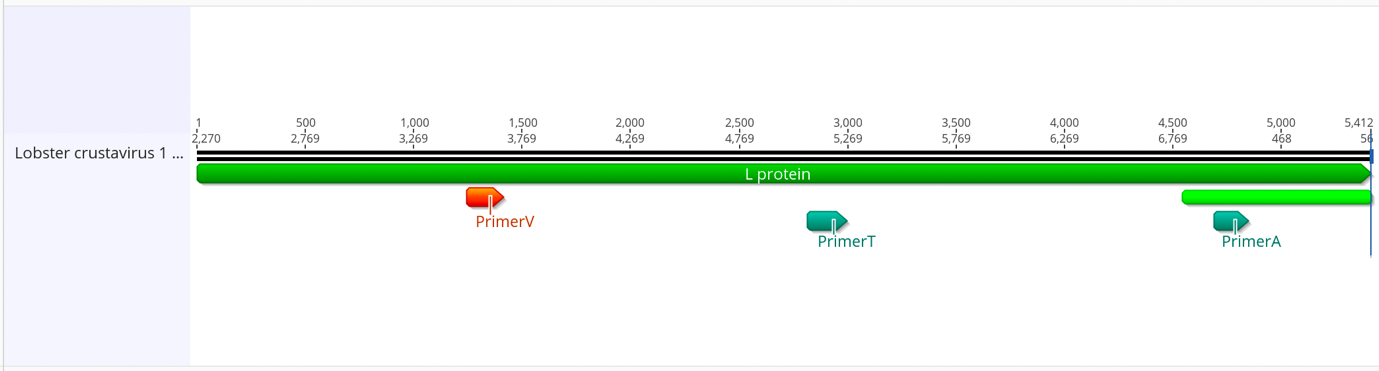

**Supplementary Figure 4.** Positions of primer set A, V, and T products across the RRLV L protein.

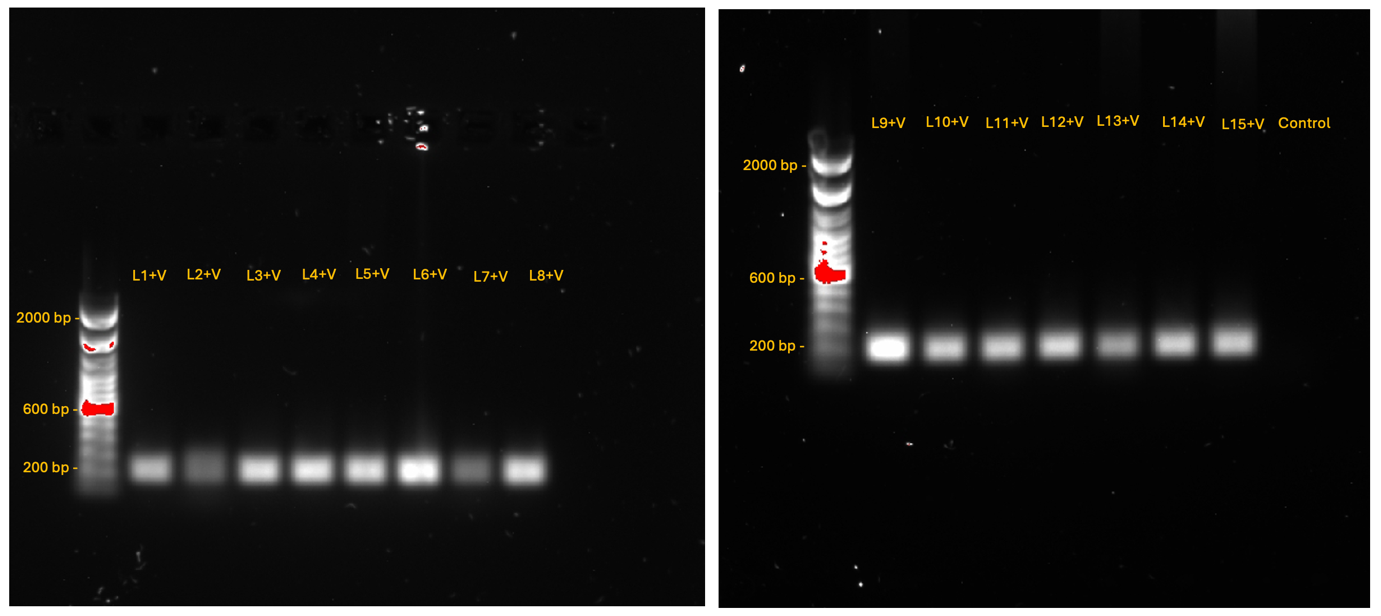

**Supplementary Figure 5** PCR confirmation of RRLCV in the haemolymph of lobsters 1-15 with primer set V.

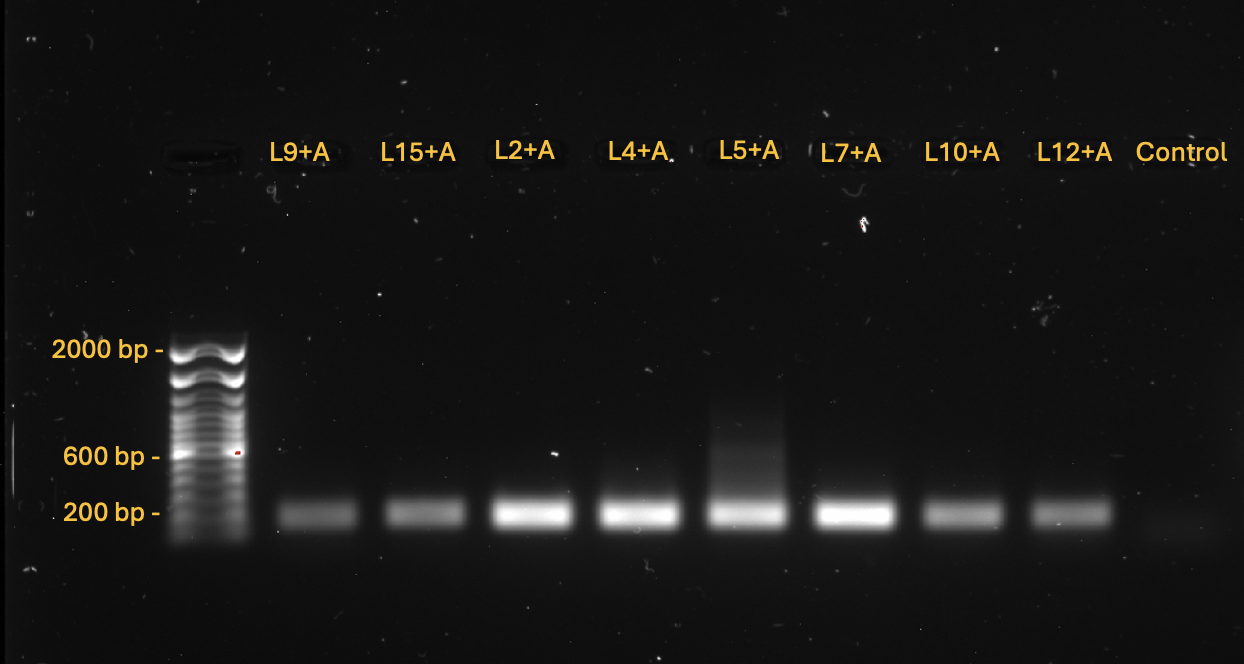

**Supplementary Figure 6.** PCR confirmation of RRLCV in the uropods of lobsters 9, 15, 2, 4, 5, 7, 10, and 12 with primer set A.

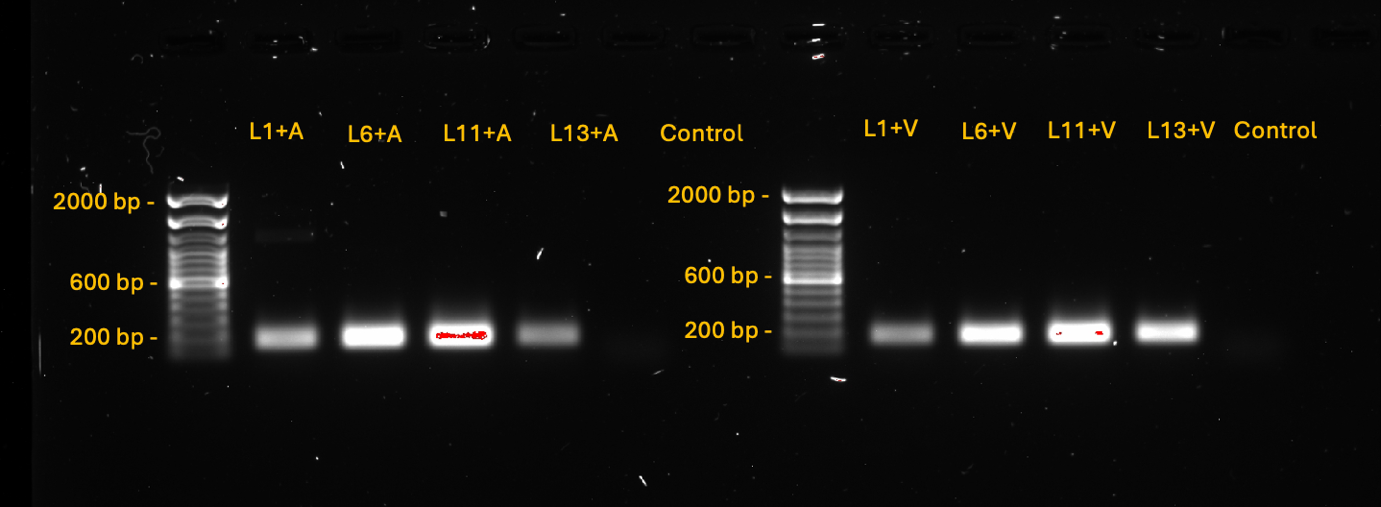

**Supplementary Figure 7.** PCR confirmation of RRLCV in the uropods of lobsters 1, 6, 11, and 12 with primer sets A (left) and V (right).

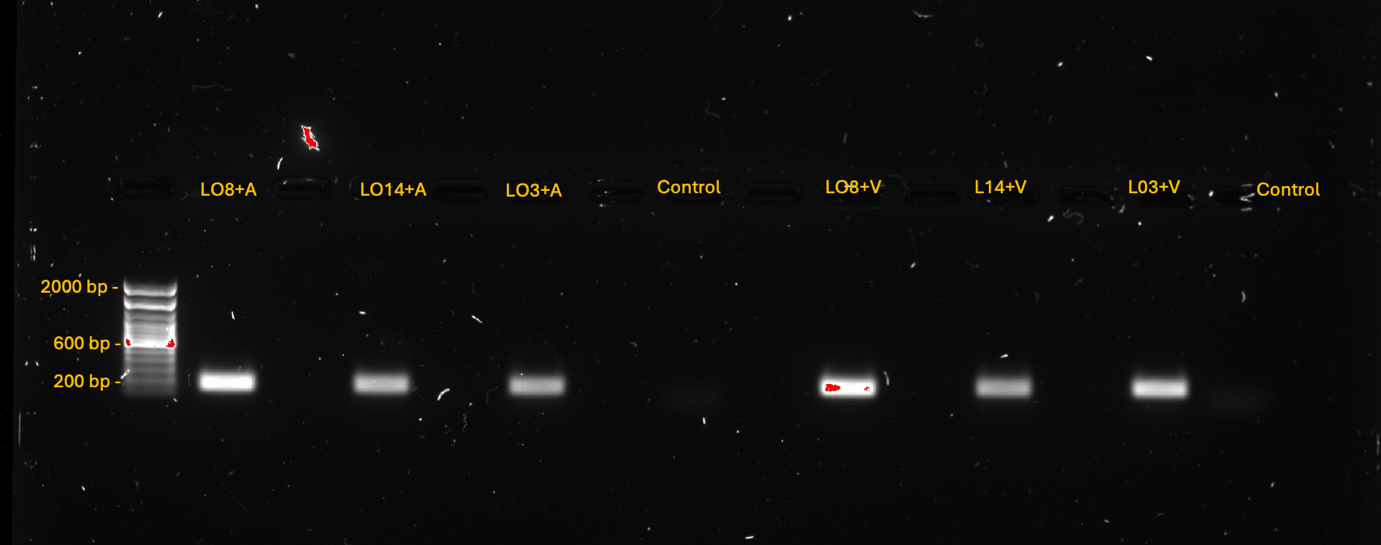

**Supplementary Figure 8.** PCR confirmation of RRLCV in the uropods of lobsters 8, 14, and 3 with primer sets A (left) and V (right).
